# Anti-trypanosomal activity and target validation of the M17 leucyl-aminopeptidase inhibitor DDD00057570 in *Trypanosoma cruzi*

**DOI:** 10.64898/2026.10.09.757876

**Authors:** Mirtha Elisa Aguado, Janaina de Freitas Nascimento, Mayke Bezerra Alencar, Maikel Izquierdo, Jorge González-Bacerio, Ariel Mariano Silber

## Abstract

**Background:** Chagas disease, caused by the parasite *Trypanosoma cruzi*, affects six to seven million people, and its treatment still relies on two drugs introduced more than fifty years ago, both of limited efficacy and poorly tolerated. The major acidic M17 leucyl-aminopeptidase of *T. cruzi* is expressed in all developmental stages of the parasite and has been proposed as a drug target, but it has not been pharmacologically validated.

**Methodology/Principal Findings:** We assessed DDD00057570, a compound originally identified as an inhibitor of the orthologue enzyme of *Leishmania major*, against the recombinant *T. cruzi* enzyme and against the parasite. The compound inhibited the recombinant enzyme with an IC_50_ of 2.7 ± 0.9 µM and reduced the proliferation of insect-stage epimastigotes, with a half-maximal effective concentration of 3.7 ± 1.1 µM. Parasites engineered to overexpress the enzyme were twice as resistant to the compound as wild-type parasites, indicating that the enzyme is a primary target in the living parasite. Treatment increased the proportion of epimastigotes with fragmented DNA. In infected mammalian cells, the compound reduced the release of trypomastigotes, the stage responsible for dissemination in the mammalian host, with a half-maximal effective concentration of 1.3 ± 0.6 µM, while its half-maximal cytotoxic concentration on CHO-K_1_ mammalian host cells was 41.5 ± 3.9 µM, resulting in a selectivity index of approximately 32.

**Conclusions/Significance:** These results validate the major M17 leucyl-aminopeptidase as a drug target in *T. cruzi* through target overexpression, and identify DDD00057570 as a selective lead structure warranting mechanistic, pharmacokinetic and *in vivo* studies toward a new treatment for Chagas disease.

**AUTHOR SUMMARY:** Chagas disease is caused by *Trypanosoma cruzi*, a parasite spread by blood-feeding triatomine insects that affects millions of people, mostly in Latin America. The only two drugs available were discovered more than half a century ago; both often fail and cause side effects severe enough that many patients abandon treatment. New drugs are urgently needed, and finding them starts with knowing which parasite proteins are worth attacking. Herein we studied an enzyme the parasite uses to trim fragments from proteins and release leucine, a building block it needs to proliferate and sustain the infection. We tested a compound originally designed to block the same enzyme in the related parasite causing leishmaniasis. We found that it blocks the recombinant enzyme, slows the proliferation of the parasite, and reduces the number of parasites bursting from infected mammalian cells, the step by which infection spreads through the body. To confirm the enzyme is what the compound attacks, we engineered parasites carrying extra copies of it; these became more resistant, exactly as expected. The compound was far more toxic to the parasite than to mammalian cells. We therefore propose this enzyme as a genuine drug target, and this compound as a starting point.

## INTRODUCTION

Chagas disease is caused by *Trypanosoma cruzi*, an euglenozoan parasite belonging to the group Kinetoplastina [1]. Current estimates indicate that between six and seven million people are infected with this parasite, with 30,000 to 40,000 new cases and approximately 12,000 deaths reported annually [2]. *T. cruzi* has a complex life cycle that alternates between reduviid insect vectors and mammalian hosts, including humans. The parasite presents four main developmental stages: epimastigotes and metacyclic trypomastigotes in the insect vector, and cell-derived trypomastigotes and amastigotes in the mammalian host. Currently, only two drugs-nifurtimox and benznidazole-are available for treatment. These nitroheterocyclic compounds require prolonged administration but are poorly tolerated due to their toxicity. As a result, there is an urgent need to develop new treatment options [3].

In the context of drug discovery for infectious diseases, a rational approach often relies on the identification, characterization, and validation of novel molecular targets. This strategy enables structure-based drug design, thereby accelerating progress in drug development programs [4, 5]. Proteases have emerged as promising therapeutic targets in a wide range of diseases [6], including those of infectious origins [7]. These enzymes play key roles in several aspects of parasite physiology and virulence, such as cell proliferation, differentiation, tissue dissemination, and infection of mammalian cells [8, 9]. Remarkably, all life cycle stages of *T. cruzi* express an acidic metallo-aminopeptidase belonging to the M17 family, which accounts for the main leucyl-aminopeptidase (LAP) activity detected in parasite extracts [10].

Amino acids have been shown to serve as important carbon and energy sources across the developmental stages of *T. cruzi* [11], influencing key processes such as infection, differentiation, and resistance to nutritional stress [12–16]. Although *T. cruzi* LAP (*Tc*LAP) has not yet been formally validated as a therapeutic target, several lines of evidence suggest that it may play important roles in the parasités survival in its hosts [10, 17, 18]. Other work showed that the *Tc*LAP inhibitor arphamenine A [19] was able to inhibit the *in vitro* proliferation of *Trypanosoma brucei brucei* [20], indicating that the relevance of LAP spans beyond *T. cruzi*. Moreover, knockdown of *Trypanosoma brucei* LAP1 (*Tb*LAP1)—an M17 LAP from *T. brucei*—results in delayed cytokinesis [21], and bestatin, a classical metallo-aminopeptidase inhibitor [22], has been reported to inhibit *Tc*LAP *in situ* in epimastigotes [18]. Notably, a potent and selective *Tc*LAP inhibitor has recently been shown to be active on intracellular *T. cruzi* amastigotes [17]. Together, these findings support the notion that *Tc*LAP inhibition represents a promising strategy to block parasite proliferation and to guide the development of novel anti-Chagas disease drugs.

Among recently identified LAP inhibitors, DDD00057570 (Fig. 1) has emerged as a promising lead candidate against leishmaniasis. It was selected for its inhibitory activity against *Leishmania major* M17 LAP (*Lm*LAP) [23] and its low cytotoxicity on human and murine macrophages. These findings not only validated *Lm*LAP as a therapeutic target in *Leishmania* spp., but also provided a rationale for investigating its potential against *T. cruzi*. In the present study, we investigated the anti-*T. cruzi* activity of DDD00057570 by assessing its inhibitory effect on *Tc*LAP and its antiparasitic activity against *T. cruzi* epimastigotes and trypomastigotes released from infected mammalian cells. Our results indicate that DDD00057570 represents a valuable starting point for the development of novel anti-chagasic drugs, supporting further investigation into its therapeutic potential and mechanism of action.

**Figure 1.**
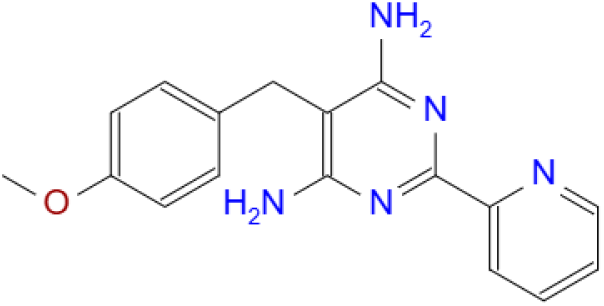
Structure of DDD00057570, originally reported as an *Lm*LAP inhibitor [23].

## 2. MATERIALS AND METHODS

### Cell culture

*T. cruzi* epimastigotes (strain CL, clone 14) [24] were maintained in the exponential proliferation phase by subculturing every 48 h in liver infusion tryptose (LIT) medium (5 g L⁻¹ liver infusion broth, 5 g L⁻¹ tryptose, 4 g L⁻¹ NaCl, 0.4 g L⁻¹ KCl, 8 g L⁻¹ Na₂HPO₄, 10 mg L⁻¹ hemin, 2 g L⁻¹ glucose, pH 7.4) [25] at 28 °C. Chinese Hamster Ovary cells (CHO-K_1_) were cultivated in RPMI-1640 medium supplemented with 10% heat-inactivated fetal calf serum (FCS), 100 µg/mL penicillin, and 100 µg/mL streptomycin at 37 °C in a humidified atmosphere containing 5% CO₂. Trypomastigotes were obtained by infection of CHO-K_1_ cells with trypomastigotes as described previously [26] and collected from the extracellular medium five to six days after infection.

### Dose-response analysis inhibition of TcLAP

Recombinant *Tc*LAP was generated as previously described [19]. Aminopeptidase activity was measured using the fluorogenic substrate Leu-7-amido-4-methylcoumarin (Leu-AMC) (Sigma-Aldrich, USA). Leu-AMC was solubilized in dimethyl sulfoxide (DMSO) (analytical grade, Sigma-Aldrich, USA) and used at a concentration corresponding to its calculated *K*◻ (1 mM) [19]. AMC release (excitation: 380 nm, emission: 460 nm) was monitored in continuous mode for 10 min using a microplate spectrofluorometer (PHERAstar FSX, BMG LABTECH, UK). Assays were performed in enzymatic activity buffer (50 mM Tris-HCl pH 9.0, 4 mM CoCl₂) at 50 °C, the optimal temperature of the recombinant enzyme [19]. The enzyme concentration (9.09 × 10⁻⁷ M, determined by the bicinchoninic acid method) was selected from the linear range of initial velocity (v₀) vs. enzyme concentration. Reactions were carried out in a final volume of 200 µL in 96-well black plates. Reaction velocities were obtained from the linear range of the progress curves, corresponding to substrate consumption below 5% (v₀ conditions). Slopes obtained from linear fits with R² < 0.98 were excluded. All assays were carried out in triplicate. For inhibition assays, *Tc*LAP was incubated with nine concentrations (0.1–100 µM range) of compound DDD00057570 (Molport, Latvia) in the enzymatic activity buffer. The mixture was preincubated for 30 min at 50 °C prior to substrate addition. DDD00057570 was solubilized in DMSO, and the final DMSO concentration did not exceed 2% (v/v). Control reactions were performed by preincubation of the enzyme with the solvent alone. Residual activity was defined as the ratio of the reaction velocity in the presence of DDD00057570 to that of the control. IC₅₀ values were calculated by nonlinear regression using the logistic function:

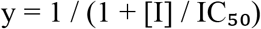

where y is residual aminopeptidase activity and [I] is inhibitor concentration. Fitting was performed with GraphPad Prism (version 8.0.2).

### Inhibitory effect of compound DDD00057570 on T. cruzi epimastigotes proliferation

The effect of the compound on *T. cruzi* epimastigote proliferation was determined through a dose–response assay. Epimastigotes in the exponential proliferation phase (2 × 10⁶ cells/mL) were seeded into 96-well culture plates (200 µL/well) and incubated at 28 °C in the presence of increasing concentrations of compound DDD00057570, ranging from 0.005 to 10 µM. Parasite growth was monitored daily for 10 days by measuring optical density (OD) at 620 nm. OD values were converted to cell density (cells/mL) using a standard calibration curve established from known parasite densities [27]. The half-maximal effective concentration (EC₅₀) was determined from cell density data collected on day 4 of proliferation, corresponding to the mid-exponential growth phase. Data were analyzed by nonlinear regression fit to a sigmoidal dose–response curve using GraphPad Prism version 8. Benznidazole (dissolved in DMSO) at a final concentration of 20 µM was used as a positive control, while untreated parasites grown in the presence of the same concentration of DMSO served as the negative control [28]. Each compound concentration was tested in quadruplicate per experiment, and the results represent three independent biological replicates.

### Validation of TcLAP as a target through its overexpression in epimastigotes

To validate *Tc*LAP as the primary target of compound DDD00057570, the sensitivity of parasites overexpressing the putative target was compared with that of parasites not overexpressing the protein. For this purpose, two transgenic *T. cruzi* lineages were generated: one stably transfected with an expression vector harboring the gene encoding *Tc*LAP (*Tc*LAP-overexpressing lineage), and a wild-type lineage.

### Cloning of the TcLAP Gene in the pTEX Expression Vector

The *TcLAP* gene (C4B63_193g9 metallo-peptidase, Clan MF, Family M17) was amplified from *T. cruzi* genomic DNA using the following primers: M005 (Fwd CDS M17 *BamHI*: AGAGGATCCATGAACAGACCTCCTGCTAC) and M006 (Rev CDS M17 *HindIII*: AGAAGCTTTTATCGTAAATTACGAAGAT). The PCR product was purified using the QIAquick PCR Purification Kit (Qiagen, MD, USA) and cloned into the pTEX vector [29] using the *BamHI* and *HindIII* restriction sites. For cloning, the purified *TcLAP* amplicon and the pTEX plasmid were independently digested with *BamHI* and *HindIII* FastDigest enzymes (Thermo Fisher Scientific, MA, USA) for 5 min at 37 °C. The digested plasmid was additionally treated with FastAP Thermosensitive Alkaline Phosphatase (Thermo Fisher Scientific, MA, USA) to prevent self-ligation. The digested insert and vector were ligated overnight at 16 °C using T4 DNA ligase (Thermo Fisher Scientific, MA, USA). The resulting pTEX-*TcLAP* construct was transformed into *E. coli* DH5α competent cells, and transformants were selected on LB agar plates containing ampicillin (100 µg/mL). Individual colonies were screened for the presence of the insert by colony PCR using primers M005 and M006, with colonies added directly to the PCR mix as the DNA template. Positive clones were cultured in liquid medium, and the recombinant plasmid was purified using the QIAprep Spin Miniprep Kit (Qiagen, MD, USA). The integrity and correct insertion of the *TcLAP* gene were confirmed by restriction enzyme digestion followed by agarose gel electrophoresis.

### Generation of T. cruzi epimastigotes lineages overexpressing TcLAP

Exponentially growing *T. cruzi* epimastigotes were arrested in the G₁/S phase by treatment with 20 mM hydroxyurea for 18 h. Parasites were then washed once with 1 mL of transfection buffer (90 mM sodium phosphate, 5 mM KCl, 0.15 mM CaCl₂, 50 mM HEPES, pH 7.2), and 4 × 10⁷ parasites were resuspended in 80 µL of the same buffer. Prior to transfection, 25 µg of the pTEX-*TcLAP* plasmid was precipitated with 0.3 M potassium acetate and 2.5 volumes of isopropanol for 18 h at –20 °C. The parasite suspension was transferred to a 0.2 cm cuvette, and 10 µL of the precipitated DNA was added. Transfection was performed using a Nucleofector 2b device (Lonza, Basel, Switzerland) set to program U-033, applying a single pulse. Immediately after electroporation, 1 mL of LIT medium supplemented with 20% FCS was added to the cuvette. The contents were then transferred to 4 mL of LIT medium with 20% FCS and incubated for 24 h at 28 °C with 5% CO₂. Following the recovery period, parasites were diluted in a 24-well plate containing LIT medium supplemented with 20% FCS, penicillin–streptomycin, and 2.5 µg/mL puromycin for selection of stable transfectant lineages. Cultures were maintained at 28 °C with 5% CO₂.

### Analysis of TcLAP Overexpression

Overexpression of *TcLAP* in transgenic epimastigotes was confirmed by assessing enzymatic activity toward the fluorogenic substrate Leu-AMC as described above, in intact parasites, using wild-type parasites as a reference. Assays were performed with 10 µM Leu-AMC in a final reaction volume of 200 µL, using PBS as the assay buffer. A normalized number of 5 × 10⁶ parasites, counted using a Neubauer chamber, were incubated with the substrate for 10 min at 37 °C. Enzymatic activity was expressed as fluorescence units per minute (FU/min) for each parasite population, calculated from the slope of the fluorescence curve generated by substrate degradation over time. Fluorescence measurements were acquired using a SpectraMax i3 plate fluorimeter (Molecular Devices, CA, USA) with excitation and emission wavelengths set to 380 nm and 460 nm, respectively. Three independent experiments were performed, each with technical quadruplicates.

### Sensitivity to DDD00057570 of parasites overexpressing TcLAP

To validate *Tc*LAP as the primary target of compound DDD00057570, concentration–response assays were performed comparing the sensitivity of *TcLAP*-overexpressing and - non-overexpressing parasites. Both strains were incubated with increasing concentrations of the compound (0–10 µM), and parasite proliferation capacity was assessed to determine the EC₅₀ for each strain. A shift in EC₅₀ values—specifically, reduced sensitivity (i.e., higher EC₅₀) in the overexpressing lineage relative to non-overexpressing parasites—would be indicative of *Tc*LAP being the primary target. Three independent experiments were performed, each with technical quadruplicates.

### Cell Cycle Analysis

Epimastigotes were cultured in LIT medium and treated with the compound at the EC₅₀ concentration for 48 h. Following treatment, parasites (1 × 10⁷ cells/mL) were collected by centrifugation at 2,700 × *g* for 5 min, washed with PBS, and fixed in 70% ethanol for 4 h. After fixation, the cells were washed twice with PBS and incubated with 10 μg/mL RNase A (Thermo Fisher Scientific, MA, USA) for 30 min at 37 °C. DNA content was assessed by staining with 20 μg/mL propidium iodide (Molecular Probes/Invitrogen, OR, USA), and samples were analyzed by flow cytometry using a BD Accuri™ C6 Plus instrument. A total of 10,000 events were collected per sample. Three biologically independent experiments were performed, each with four technical replicates. Histograms and gates for each cell cycle phase were analyzed using BD CSamplerPlus Software (v1.0.27.1) and FlowJo software (v10.8.1), with cell cycle distribution fitted using the model incorporated in FlowJo.

### Effect of the Compound on Trypomastigote Bursting

For infection assays, CHO-K_1_ cells seeded in 24-well plates were infected with *T. cruzi* trypomastigotes at a multiplicity of infection (MOI) of 50 (2.5 × 10⁶ parasites per well) for 3 h. After infection, the supernatant containing non-internalized parasites was removed, and monolayers were washed twice with PBS. Fresh RPMI-1640 medium supplemented with 10% FCS was added, and infected cells were treated daily with the compound at concentrations ranging from 0.1 to 10 µM from day 0 to day 4 post-infection. Cultures were maintained at 37 °C with 5% CO₂ for the first 24 h, after which the medium was replaced with RPMI-1640 containing 2% FCS, and cultures were shifted to 33 °C with 5% CO₂. On days 4, 5, 6, and 7 post-infections, the number of trypomastigotes released into the supernatant was quantified using a Neubauer hemocytometer. All measurements were performed in three independent experiments, each with quadruplicate technical replicates.

### Viability Assay in CHO-K_1_ Cells

CHO-K_1_ cells were seeded in 96-well plates at a density of 1.0 × 10⁵ cells per well in 100 µL of RPMI-1640 medium supplemented with 10% FCS. Following a 24 h incubation to allow cell adhesion, cells were exposed to increasing concentrations of the compound for 96 h. Untreated cells served as negative controls. Cell viability was assessed using the MTT (3-(4,5-dimethylthiazol-2-yl)-2,5-diphenyltetrazolium bromide) assay, as previously described [30]. The formazan crystals produced by viable cells were solubilized in DMSO, and the optical density (OD) was measured at 540 nm with a reference wavelength of 690 nm. The half-maximal cytotoxic concentration (CC₅₀) was determined by fitting the data to a sigmoidal dose–response curve using GraphPad Prism version 8. Three independent experiments were performed, each with quadruplicate technical replicates.

### Statistical Analysis

All dose-response analyses and statistical tests were conducted using GraphPad Prism v10.1.0. Histograms for each cell cycle phase were analyzed using BD CSamplerPlus Software (v1.0.27.1) and FlowJo software (v10.8.1), with cell cycle distribution fitted using the model incorporated in FlowJo. For comparisons between paired experiments we used the t-test while for comparisons of multi-sample experiments were made by using a 2-way ANOVA followed by the Tuckey post-test. All data (except those for the enzyme inhibition curve) are presented as mean ± SEM from three independent experiments, each performed in quadruplicate.

## RESULTS

### DDD00057570 inhibits the proliferation of T. cruzi epimastigotes

To initially investigate the trypanocidal activity of DDD00057570, its ability to inhibit the *in vitro* proliferation of *T. cruzi* epimastigotes was assessed. Benznidazole, the reference drug for Chagas disease, was used as a positive control to validate the assay (Fig. 2A). Under these conditions, DDD00057570 demonstrated trypanocidal activity, with an EC_50_ value of 3.7± 1.1 μM for inhibition of epimastigote proliferation after 96 hours of treatment (Fig. 2B).

**Figure 2.**
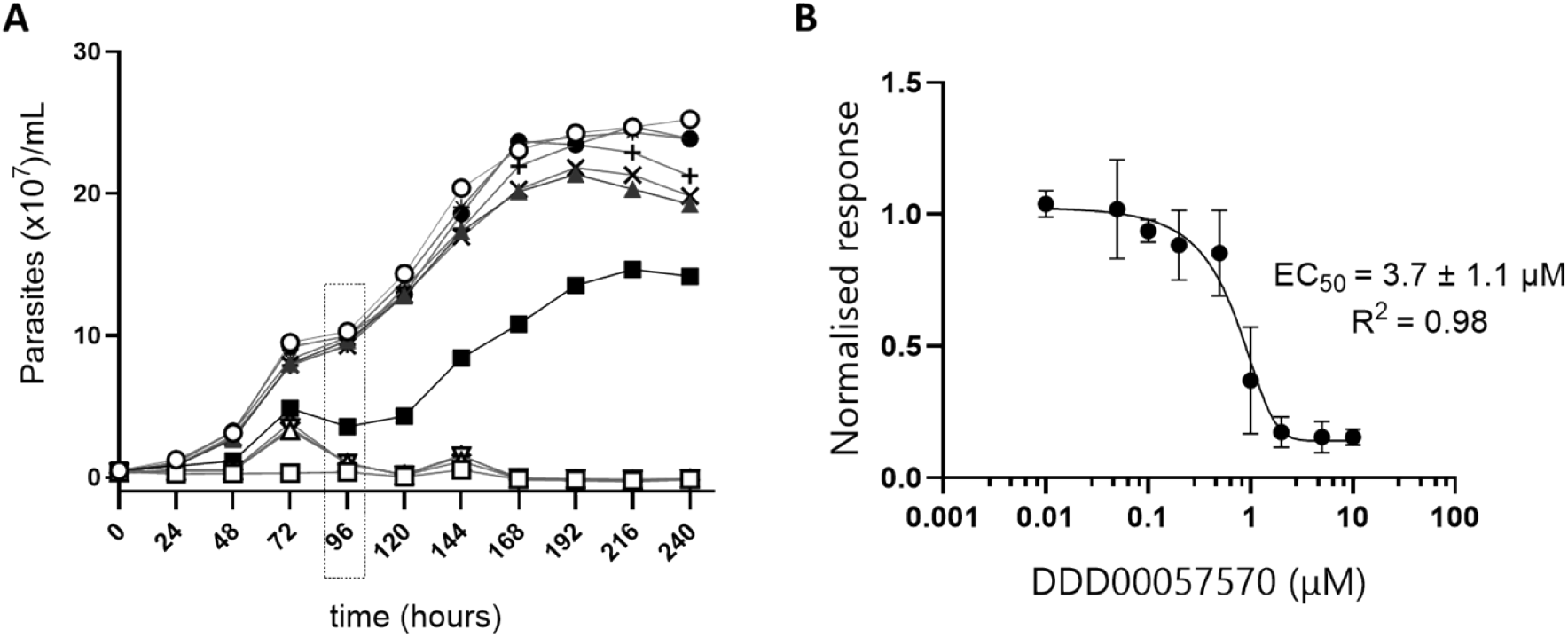
*In vitro* anti-*T. cruzi* activity of DDD00057570 against epimastigotes. (A) Proliferation curves of *T. cruzi* epimastigotes treated with increasing concentrations of DDD00057570. The symbols represent: (Δ) 10 µM, (▽) 5 µM, (◊) 2.5 µM, (▪) 1.25 µM, (▴) 0.6 µM, (x) 0.3 µM, (+) 0.15 µM, (*) 0.05 µM, (●) 0.02 µM, (○) DMSO as negative control and (**□**) Benznidazole 20 μM as positive control. (B) Dose–response curve based on cell density measurements at 96 h across different concentrations of DDD00057570. Data are presented as mean ± SEM from three independent experiments, each performed in quadruplicate.

### TcLAP is a molecular target of DDD00057570 in T. cruzi

Given that DDD00057570 was originally identified as an inhibitor of *L. major* M17 leucine aminopeptidase (*Lm*LAP) [23], its inhibitory activity was assessed against the recombinant ortholog from *T. cruzi*, *Tc*LAP (Fig. 3). The compound acted as a relatively potent inhibitor of *Tc*LAP, with an IC_50_ value of 2.7 ± 0.9 μM.

**Figure 3.**
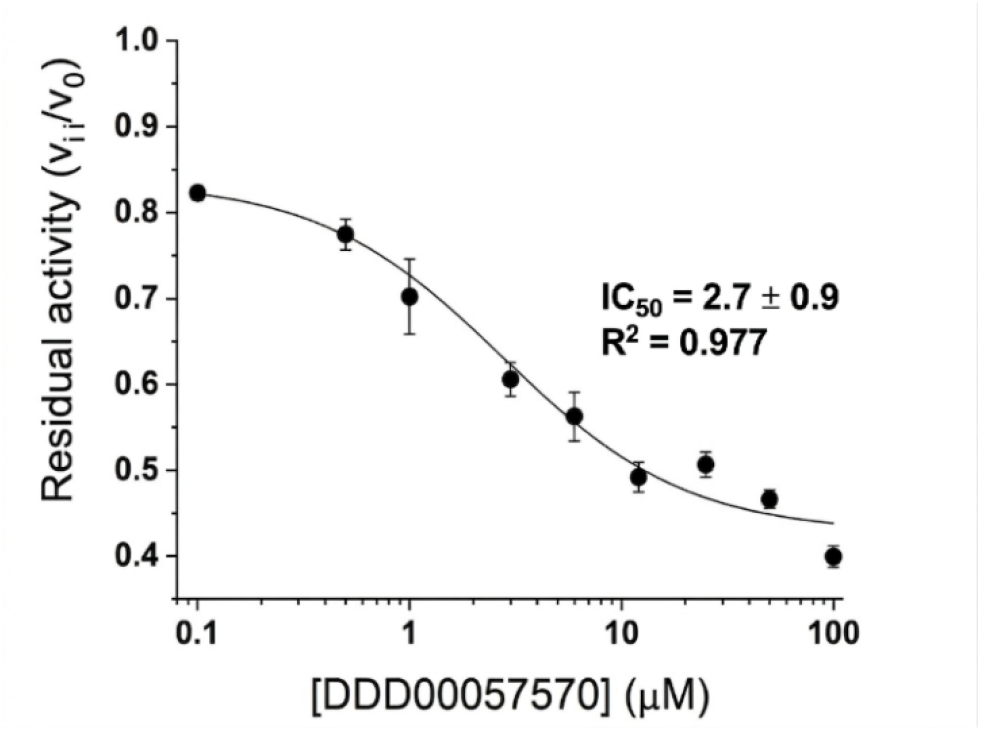
Concentration–response curve for inhibition of recombinant *Tc*LAP by DDD00057570. A representative curve from three independent biological experiments is shown. Data are presented as mean ± SD (n = 3 per concentration). The IC_50_ value is reported as mean ± SEM, along with the coefficient of determination (R²).

To confirm that *Tc*LAP is the main cellular target of DDD00057570 in *T. cruzi*, epimastigotes were transfected with the pTEX-*TcLAP* construct or with the wild-type as a control, allowing for overexpression of the putative target enzyme or maintenance of basal expression levels, respectively. Successful overexpression was confirmed by measuring LAP enzymatic activity (Fig. 4A). Lineage 4, which exhibited the highest LAP activity (Fig. 4A), was selected for this analysis. Importantly, the overexpression of *TcLAP* produced a slight diminution in the proliferation of the overexpressing lineage when compared to the wild type. The sensitivity of *Tc*LAP-overexpressing epimastigotes to the inhibitor was compared with that of wild-type parasites expressing basal levels of the enzyme. Following 96 h of exposure, *TcLAP*-over-expressing parasites showed increased resistance to DDD00057570 (Fig. 4C and D) compared to wild-type, showing an EC_50_ of 6.0 ± 0.7 *vs.*3.0 ± 1.1 μM, respectively (Fig. 4E).

**Figure 4.**
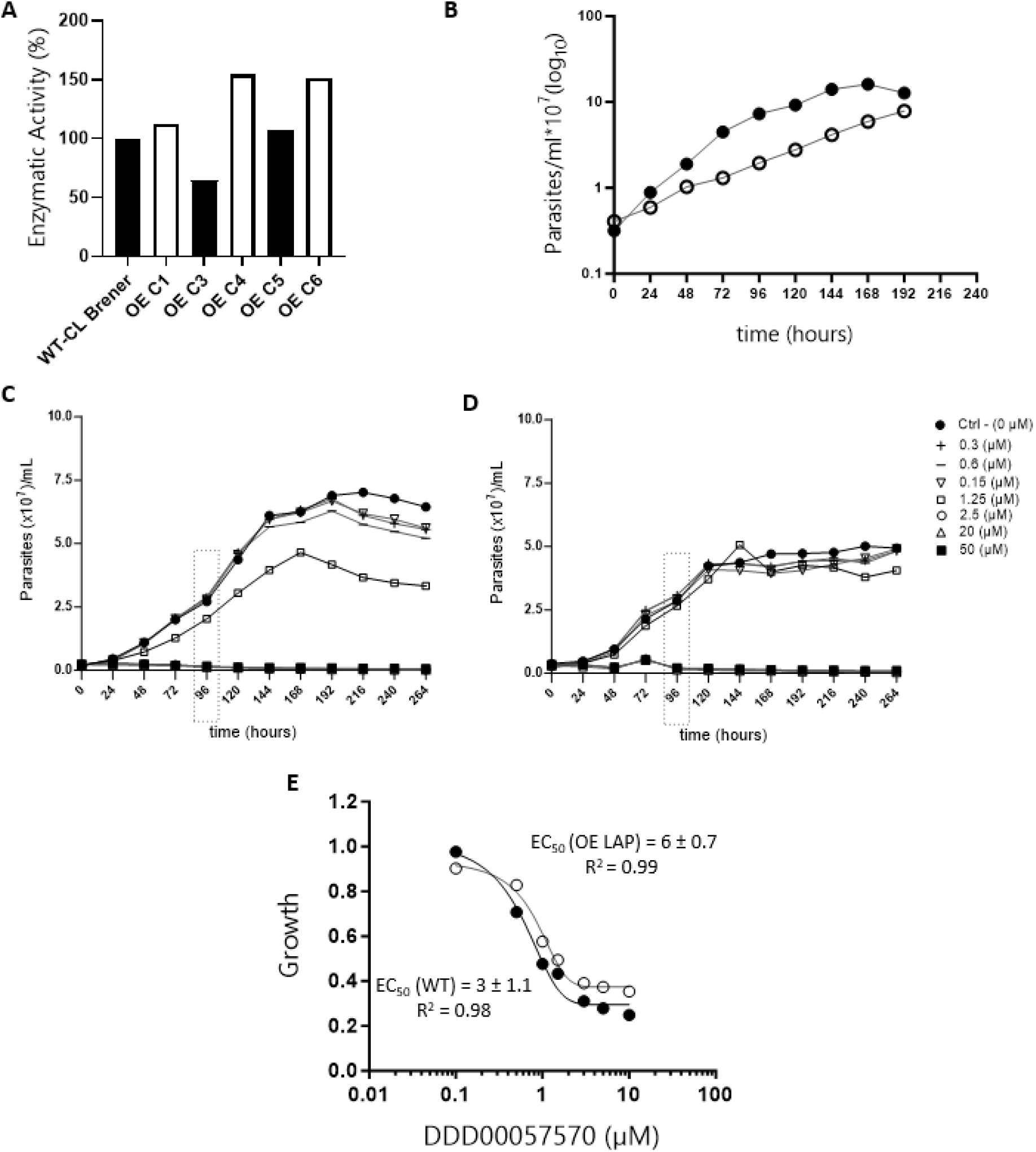
Validation of *Tc*LAP as a target in *Trypanosoma cruzi*. (A) LAP enzymatic activities of five clones of transgenic *Tc*LAP-overexpressing and wild-type epimastigotes. FU: fluorescence units. (B) Proliferation curves for the *Tc*LAP-overexpressing (Clone 4: o) and wild-type epimastigotes (●). (C) and (D) Concentration-response growth curves for the *in vitro* anti-epimastigote activity of DDD00057570 on wild-type and transgenic *Tc*LAP-overexpressing, respectively. (E) Concentration-response curves for the *in vitro* anti-chagasic activity of DDD00057570 on transgenic *Tc*LAP-overexpressing (Clone 4: o) and wild-type epimastigotes (●) at 96 h of exposure. Data represent means ± SEM from three independent experiments, each performed in quadruplicate.

### DDD00057570 alters the cell cycle of T. cruzi

The inhibitory effect of DDD00057570 on parasite proliferation may result from direct trypanocidal activity or from interference with the cell cycle, as previously reported for *Tb*LAP1 [21]. To investigate this, the cell cycle distribution of epimastigotes treated with the compound was analyzed by flow cytometry. Treatment with DDD00057570 at its EC_50_ concentration for 48 h resulted in a significant increased sub-G1 population (18.6 ± 1.2%) compared to untreated controls (8.1 ± 2%) (Fig. 5A), suggesting DNA fragmentation, consistent with cell death processes. Furthermore, the reduction in the G1/S population (treated: 28 ± 4.1 % vs control: 34.4 ± 4.3%) and the concomitant accumulation of cells in S/G2 phases (21.3 ± 3 % vs 14.2 ± 2.6 %) are consistent with an arrest before mitotic entry (Fig. 5A and B).

**Figure 5.**
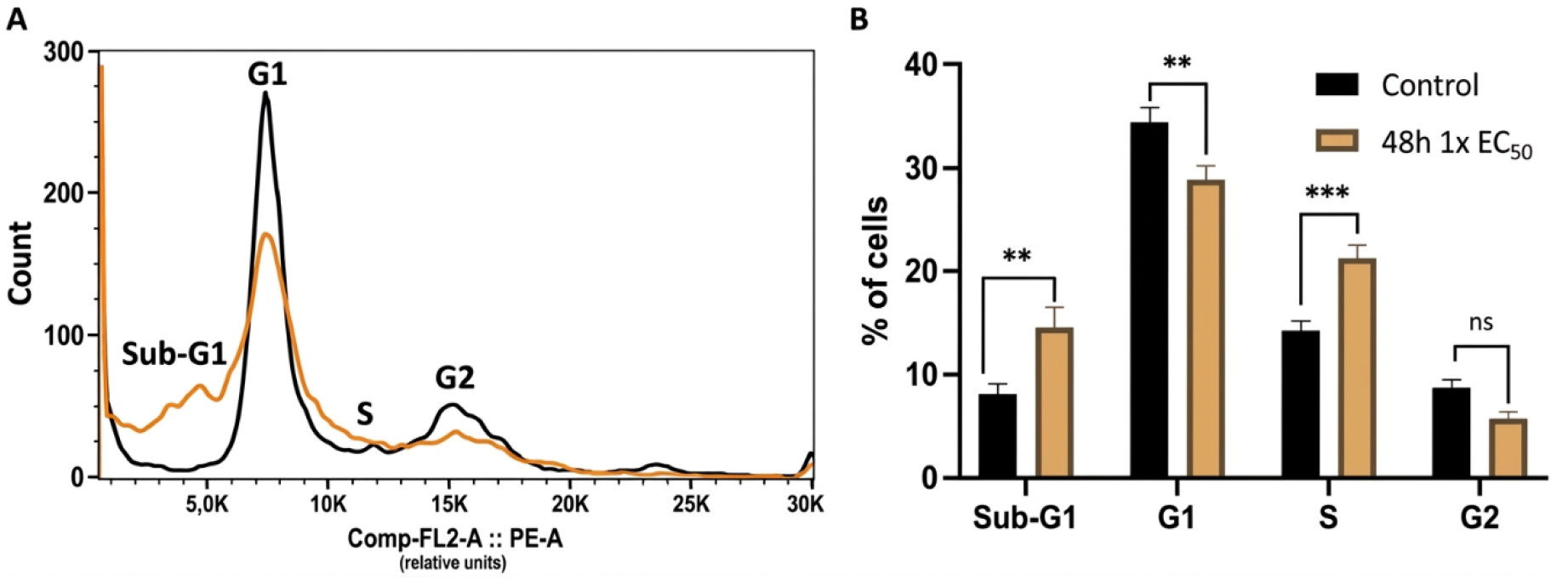
Effect of DDD00057570 on the cell cycle of *T. cruzi* epimastigotes assessed by flow cytometry. Parasites were treated with the EC50 concentration of DDD00057570 for 48 h and stained with propidium iodide for DNA content analysis. A total of 10,000 events were analyzed per sample. **(A)** Representative histograms of DNA content from one experiment. **(B)** Quantification of cells in each phase of the cell cycle. Data represent mean ± SEM from three independent experiments, each performed in quadruplicate. Statistical comparisons to the control were performed using a t-test (**, p < 0.01; ***, p < 0.001; ns, not significant).

### DDD00057570 shows relatively low cytotoxicity in mammalian CHO-K1 cells

To assay the effect of DDD00057570 in the relevant clinical stages of *T. cruzi*, the maximum non-cytotoxic concentration for the mammalian cell to be infected (in our case, CHO-K_1_) must be established. The cells were cultured until sub-confluence and maintained in the presence of different concentrations of the compound (ranging from 0.02 to 100 µM) for 96 h. The formation of formazan from MTT was used as a proxy for cell viability. Under these conditions, DDD00057570 did not exhibit significant cytotoxicity toward mammalian cells, with a maximum viability reduction of 33% observed at the highest concentration tested relative to untreated controls. A logistic dose–response non-linear regression curve was fitted to the experimental data to estimate the compound’s toxicity, yielding a CC_50_ value of 41.5 ± 3.9 μM (Fig. 6)

**Figure 6.**
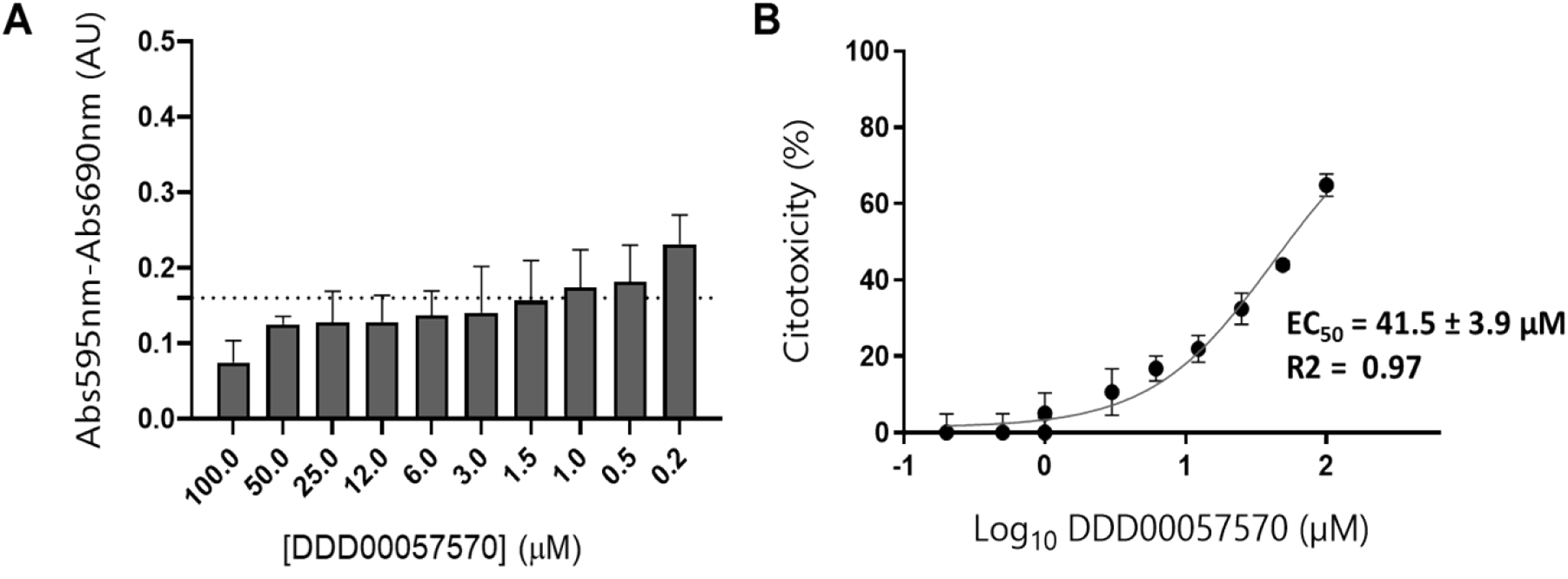
Cytotoxicity of DDD00057570 on mammalian CHO-K_1_ cells. **(A)** Viability of CHO-K1 cells treated with increasing concentrations of DDD00057570 (0–100 μM) was assessed by the MTT assay. The dotted line represents the control in the absence of the compound (DMSO only). **(B)** Concentration–response curve following 96 h of compound exposure. Data represent mean ± SEM from three independent experiments, each performed in quadruplicate.

### DDD00057570 inhibits trypomastigote egress from infected host cells

The effect of DDD00057570 on parasite infection of mammalian cells was assessed using CHO-K_1_ cells infected with cell-derived trypomastigotes. The number of trypomastigotes released from infected cells was recorded from 4 days post-infection onward. Treatment with DDD00057570 resulted in a concentration-dependent reduction in trypomastigote egress (Fig. 7A), with an EC_50_ value of 1.3 ± 0.6 μM (Fig. 7B).

**Figure 7.**
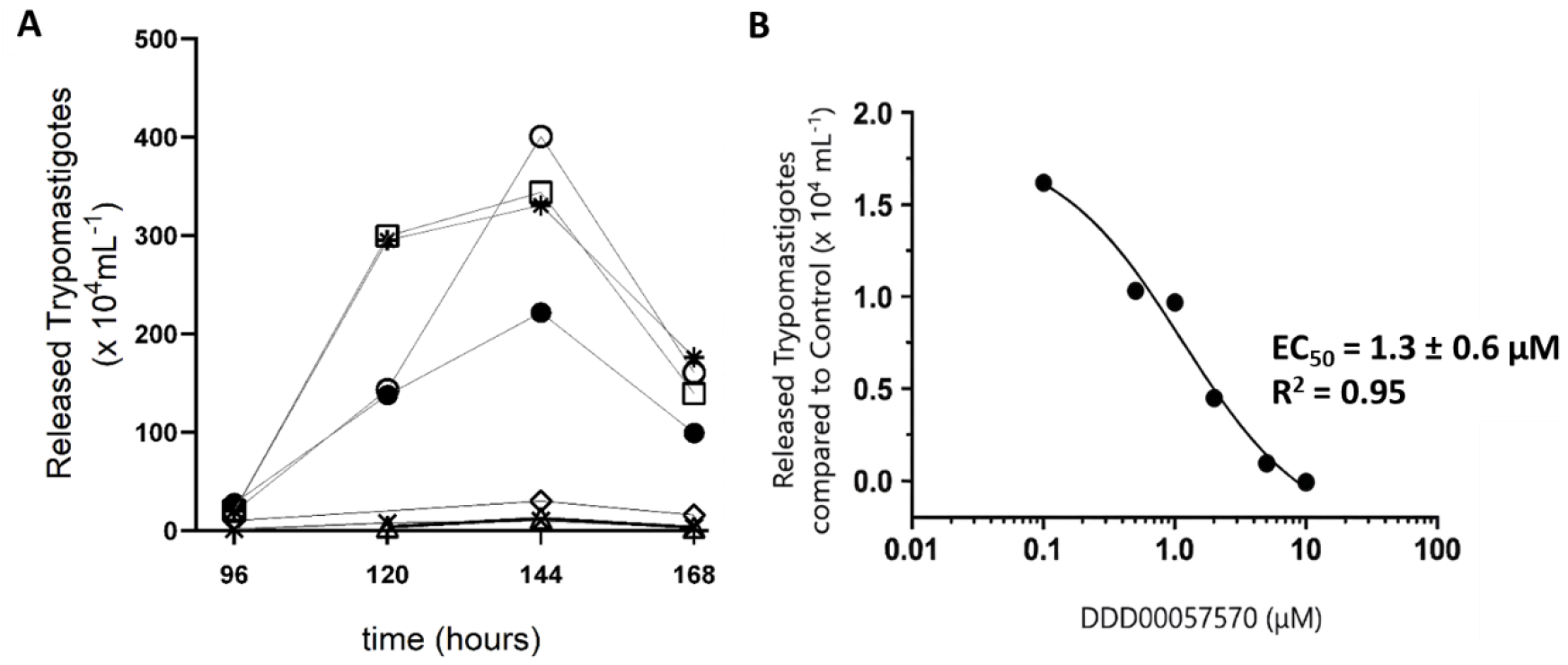
Effect of DDD00057570 on infection of CHO-K_1_ cells by *T. cruzi* trypomastigotes. **(A)** Number of trypomastigotes released from infected cells following treatment with increasing concentrations of DDD00057570, measured from day 4 post-infection onward. The symbols represent the following experimental conditions: (o) untreated control (CHO + trypomastigotes), and increasing concentrations of the compound: (Δ) 10 µM, (x) 5 µM, (◊) 2.5 µM, (□) 1.25 µM, (*) 0.5 µM and (●) 0.1 µM. **(B)** Concentration–response curve for inhibition of trypomastigote release at day 4 post-infection. Data are presented as mean ± SEM from three independent biological replicates, each performed in quadruplicate.

## 4. DISCUSSION

M17 leucine aminopeptidases (LAPs) have emerged as attractive drug targets in parasitic diseases due to their involvement in numerous essential physiological processes across diverse pathogens [10, 17, 23, 31–35]. In trypanosomatids, these enzymes contribute to critical steps such as infection establishment, proliferation, differentiation, immune evasion, and tissue dissemination [31, 36]. Despite their central roles, only a few M17 LAPs from trypanosomatids have been experimentally validated as therapeutic targets [21, 23].

Beyond trypanosomatids, M17 LAPs also hold therapeutic potential in other parasitic organisms. For instance, the M17 LAP of *Plasmodium falciparum* (PfA-M17) is indispensable for hemoglobin degradation and essential housekeeping functions, as evidenced by studies using a specific bestatin-based inhibitor [34, 37]. Similarly, in *Acanthamoeba castellanii*, reduced M17 LAP expression or chemical inhibition with agents such as 1,10-phenanthroline or bestatin causes severe ultrastructural defects and impairs encystation [38]. The antiparasitic effect of bestatin against *Babesia bovis* has also been linked to inhibition of its M17 LAP [39]. Additionally, *Schistosoma mansoni* encodes two M17 LAPs involved in hemoglobin digestion, surface membrane remodelling, and egg hatching, as demonstrated by RNAi knockdown and pharmacological inhibition [40]. Collectively, these findings position M17 LAPs as a conserved and functionally vulnerable node across disparate parasite lineages, reinforcing their potential as high-value targets for broad-spectrum antiparasitic drug development.

*TcLAP* is expressed in all life cycle stages of *T. cruzi*: epimastigotes, trypomastigotes, and amastigotes [10]. The parasite lacks the biosynthetic pathway for branched-chain amino acids and must acquire them through uptake from the extracellular environment or via proteolysis [41]. This dependency is particularly relevant under conditions of nutrient limitation, such as those encountered in the vector insect midgut [42, 43] or during competition with the metabolic pathways once in the cytoplasm of the vertebrate host cell [11]. Although *Tc*LAP participates in the release of leucine from proteins and peptides for use by the parasite’s metabolic machinery, its role in homeostasis likely extends beyond branched-chain amino acid supply. Proposed functions include participation in peptide detoxification [44], membrane protein remodeling [45], degradation of abnormal proteins [44], and osmoregulation [46]. Due to this accumulation of multiple functions, it has been proposed that *Tc*LAP could be a promising target to identify anti-chagasic agents [17].

M17 LAP inhibitors are typically small dipeptide-or tripeptide-like compounds featuring hydrophobic and bulky functional groups. To date, only a few inhibitors have been identified for *Tc*LAP, including bestatin [10], the bestatin-derived peptidomimetic KBE009 [47], arphamenine A [19], and compound 4 [17]. In contrast, a remarkably wide array of structurally diverse inhibitors has been reported for *Plasmodium falciparum* M17 LAP (PfA-M17). Beyond bestatin and nitrobestatin—both characterized by isobutyl and benzyl groups [37, 48] —PfA-M17 is also susceptible to inhibition by the bestatin-based probe Phe-Naphthyl [34], the phosphinate dipeptide analogue Co4, which contains two phenyl rings [48], and several hydroxamates: 13d (bearing pyrazole, phenyl, and *tert*-butyl groups [49]), 10o (featuring two phenyl and *tert*-butyl groups [50]), and 6k (which incorporates three bulky hydrophobic rings [51]). This disparity in inhibitor diversity highlights a critical gap in *Tc*LAP research and underscores the opportunity to expand the chemical space explored for this target.

The inhibition of *Tc*LAP by bestatin in *T. cruzi* epimastigotes demonstrates that low-molecular-weight bestatin-like compounds can engage the target intracellularly [18]. Identifying such inhibitors is especially relevant given the current treatment landscape for Chagas disease, which remains dependent on benznidazole and nifurtimox—drugs with limited efficacy and significant toxicity [2, 3]. In this context, the previously reported *Lm*LAP inhibitor DDD00057570 showed promising in vitro activity against *Leishmania* spp. [23].

Here, we evaluated both its ability to inhibit *Tc*LAP and its trypanocidal activity, extending the therapeutic potential of this compound to *T. cruzi*.

Micromolar-or submicromolar-range compounds are frequently prioritized in later stages of drug development, as they provide a suitable foundation for optimization via medicinal chemistry to enhance potency and drug-like properties [4]. The IC₅₀ value of DDD00057570 obtained here for recombinant *Tc*LAP inhibition (2.7 ± 0.9 μM) is comparable to that reported for recombinant *Lm*LAP (3.3 μM) [23]. Notably, this value is also lower than the IC₅₀ of bestatin toward *Lm*LAP (6.1 μM). Furthermore, DDD00057570 exhibits high selectivity for recombinant *Lm*LAP over recombinant human LAP, according to Aguado and co-workers [23]. To confirm the suitability of this molecule as a trypanocidal agent, its effect on epimastigote proliferation was assessed, yielding an EC₅₀ close to that reported for benznidazole (4 μM) [52].

The structural features of DDD00057570 align with those of a typical M17 LAP inhibitor. As a low-molecular-weight compound, it incorporates bulky hydrophobic groups capable of engaging the enzyme’s active site pockets through hydrophobic interactions. Key structural elements include: i) a central pyrimidine ring that imparts rigidity and limits conformational flexibility; ii) aromatic moieties (e.g., phenyl-like groups) that mimic the hydrophobic side chains of natural substrates; and iii) protonated amines (Fig. 1) that facilitate ionic interactions with negatively charged residues within the *Tc*LAP active site [34, 37, 48]. Collectively, these features are consistent with previously characterized M17 LAP inhibitors and support both membrane permeation and effective target engagement [50].

While the anti-epimastigote activity is promising, attributing it primarily to *Tc*LAP inhibition requires additional validation. Increased resistance in parasites overexpressing the aminopeptidase can be explained by two concomitant mechanisms: a larger pool of functional target protein necessitates a higher inhibitor concentration for effective blockade, and the excess target may sequester the compound, reducing its intracellular availability [53]. Collectively, the results presented in this work strongly support the conclusion that DDD00057570 exerts its *in vitro* anti-trypanosome activity through inhibition of endogenous *Tc*LAP. To better understand the functional consequences of *Tc*LAP inhibition, we investigated its effect on the parasite cell cycle. Treatment with DDD00057570 increased the sub-G1 epimastigote population, suggesting interference with DNA replication and cell division—a mechanism of action also proposed for benznidazole [54] and aromatic diamidines [55].

Having observed that the inhibitor interferes with epimastigote proliferation, we next assessed whether it could impair mammalian host cell infection or intracellular development into trypomastigotes, the forms responsible for disease dissemination in the mammalian host. Importantly, DDD00057570 exhibited low cytotoxicity against CHO-K_1_ cells, enabling evaluation across a broad range of concentrations in infected cells. This assay revealed that the compound suppresses trypomastigote release at low micromolar concentrations, which— combined with its high CC₅₀ in mammalian cells—points to remarkable selectivity, a hallmark of desirable antiparasitic agents [56]. This selectivity is further supported by its efficacy against related protozoa (e.g., *Leishmania* spp. [23]), suggesting broad applicability for kinetoplastid infections.

Despite the fact that this study identifies DDD00057570, a *Tc*LAP inhibitor, as a relevant hit for developing ant-*T. cruzi* drugs, some limitations should be acknowledged. A genetic validation of this enzyme and a more information on the role of *Tc*LAP in the biology of *T. cruzi* must be obtained by gene knockout or knockdown. A shift in EC₅₀ upon overexpression is consistent with, but does not formally prove, exclusive on-target action, and contributions from additional targets cannot be excluded. More data on the properties of DDD00057570 such as cytotoxicity assays in a variety of cell lines, physicochemical properties, metabolic stability, pharmacokinetics or *in vivo* efficacy remains to be explored. Addressing these points is the priority for the next stage of optimization of this chemical series. Collectively, our findings position DDD00057570 as a promising starting point for further optimization as an antiparasitic agent.

